# Quantitative flow cytometric selection of tau conformational nanobodies specific for pathological aggregates

**DOI:** 10.1101/2023.05.13.540640

**Authors:** Jennifer M. Zupancic, Matthew D. Smith, Hanna Trzeciakiewicz, Mary E. Skinner, Sean P. Ferris, Emily K. Makowski, Michael J. Lucas, Nikki McArthur, Ravi S. Kane, Henry L. Paulson, Peter M. Tessier

**Author notes:** To whom correspondence should be addressed: Peter M. Tessier Address: North Campus Research Complex, B10-179 2800 Plymouth Road University of Michigan Ann Arbor, MI 48109.

## Abstract

Single-domain antibodies, also known as nanobodies, are broadly important for studying the structure and conformational states of several classes of proteins, including membrane proteins, enzymes, and amyloidogenic proteins. Conformational nanobodies specific for aggregated conformations of amyloidogenic proteins are particularly needed to better target and study aggregates associated with a growing class of associated diseases, especially neurodegenerative disorders such as Alzheimer’s and Parkinson’s diseases. However, there are few reported nanobodies with both conformational and sequence specificity for amyloid aggregates, especially for large and complex proteins such as the tau protein associated with Alzheimer’s disease, due to difficulties in selecting nanobodies that bind to complex aggregated proteins. Here, we report the selection of conformational nanobodies that selectively recognize aggregated (fibrillar) tau relative to soluble (monomeric) tau. Notably, we demonstrate that these nanobodies can be directly isolated from immune libraries using quantitative flow cytometric sorting of yeast-displayed libraries against tau aggregates conjugated to quantum dots, and this process eliminates the need for secondary nanobody screening. The isolated nanobodies demonstrate conformational specificity for tau aggregates in brain samples from both transgenic tau mouse models and human tauopathies. We expect that our facile approach will be broadly useful for isolating conformational nanobodies against diverse amyloid aggregates and other complex antigens.

## 2. INTRODUCTION

The smallest antibody fragments which retain the ability to bind antigens are single-domain antibodies, often termed nanobodies (1,2). These fragments represent the variable region of heavy chain antibodies produced by camelids (2). Nanobodies have generated much interest given their many desirable properties, including their potential to recognize conformational epitopes due to their unique binding sites, which are frequently convex in nature. Antibody-and nanobody-based discrimination between different conformations of the same protein has broad impacts ranging from structural biology studies to the development of therapies for diseases associated with protein conformational changes. For instance, nanobodies have frequently been generated to selectively recognize specific conformational states of various membrane proteins, such as G-protein coupled receptors (GPCRs) (3–12) as well as transport and channel proteins (13–16), stabilizing such proteins in particular states of activation or membrane orientation and allowing for elucidation of their structures and mechanisms. Nanobodies have also been generated to stabilize enzymes in various conformations to study their structural changes and better understand their mechanisms and overall functions (17–19). Furthermore, a limited number of nanobodies have also been developed to recognize conformational states of various proteins that undergo aggregation (20–22).

However, the potential of nanobodies to target aggregated antigens is relatively unexplored due to challenges involved in working with these complex, often insoluble antigens. In particular, the aggregation of amyloidogenic proteins represents a highly active area of research, and the development of nanobodies in this area has the potential to impact the understanding of a number of diseases associated with protein aggregation, especially neurodegenerative diseases such as Alzheimer’s and Parkinson’s diseases that are rapidly growing in prevalence (23,24). Surprisingly few nanobodies have been generated with both conformational and sequence specificity for amyloidogenic aggregates (20–22), and only one has been reported for a complex amyloidogenic protein (α-synuclein, 140 amino acids) (20).

There is broad interest in developing conformational nanobodies against other complex amyloidogenic proteins, including tau, a large protein (441 amino acids for the longest isoform) associated with Alzheimer’s disease. However, to date no tau nanobodies have been reported with both conformational and sequence specificity, and only a few tau nanobodies have been reported that are sequence-specific (25–27) or phospho-specific (28). The paucity of tau conformational nanobodies can be largely explained by the limitations of the methods used previously to generate them. The majority of previously reported nanobodies specific for amyloidogenic peptides and proteins have been isolated using either immunization followed by preparation and panning of phage libraries (22,29,30) or direct panning of synthetic phage libraries (21,25,26,31). However, it is difficult to use either method, without extensive secondary screening, to routinely isolate nanobodies specific for amyloid aggregates with a combination of three desirable binding properties: i) high sequence specificity (i.e., strong preference for tau aggregates relative to non-tau aggregates); ii) high conformational specificity (i.e., strong preference for aggregates relative to monomeric protein); and iii) low off-target binding (i.e., low binding to non-tau proteins).

In this work, we have sought to address these challenges associated with generating nanobodies with both conformational and sequence specificity for amyloid aggregates formed by large and complex proteins. We reasoned that many of the previous challenges could be addressed using quantitative flow cytometric sorting of yeast-displayed libraries to enable direct selection of nanobodies that bind selectively to tau fibrils. Herein, we report the identification of tau conformational nanobodies from immune libraries with desirable combinations of binding and biophysical properties without the need for secondary screening to identify conformational nanobodies. Moreover, we demonstrate that these nanobodies are specific for pathological tau aggregates formed in both a transgenic mouse model (P301S) and human tauopathies.

## 3. RESULTS

### 3.1 Isolation of tau conformational nanobodies from llama immunization

To generate tau conformational nanobodies, we first immunized a llama with tau fibrils (see Methods for details), and after we observed an increase in tau binding signal via serum testing (**Fig. S1**), we isolated bulk lymphocytes and generated an immune nanobody library in a standard yeast display format (**Fig. 1**). We observed that immunization with fibrils formed from a truncation of full-length tau (dGAE fibrils) led to an increase in antibody binding observed in the serum to both this fragment of tau and full-length tau fibrils (HT40 fibrils). We therefore chose to perform subsequent sorting using HT40 tau fibrils with the goal of detecting conformational binding to full-length tau fibrils, which are found *in vivo*.

**Figure 1.**
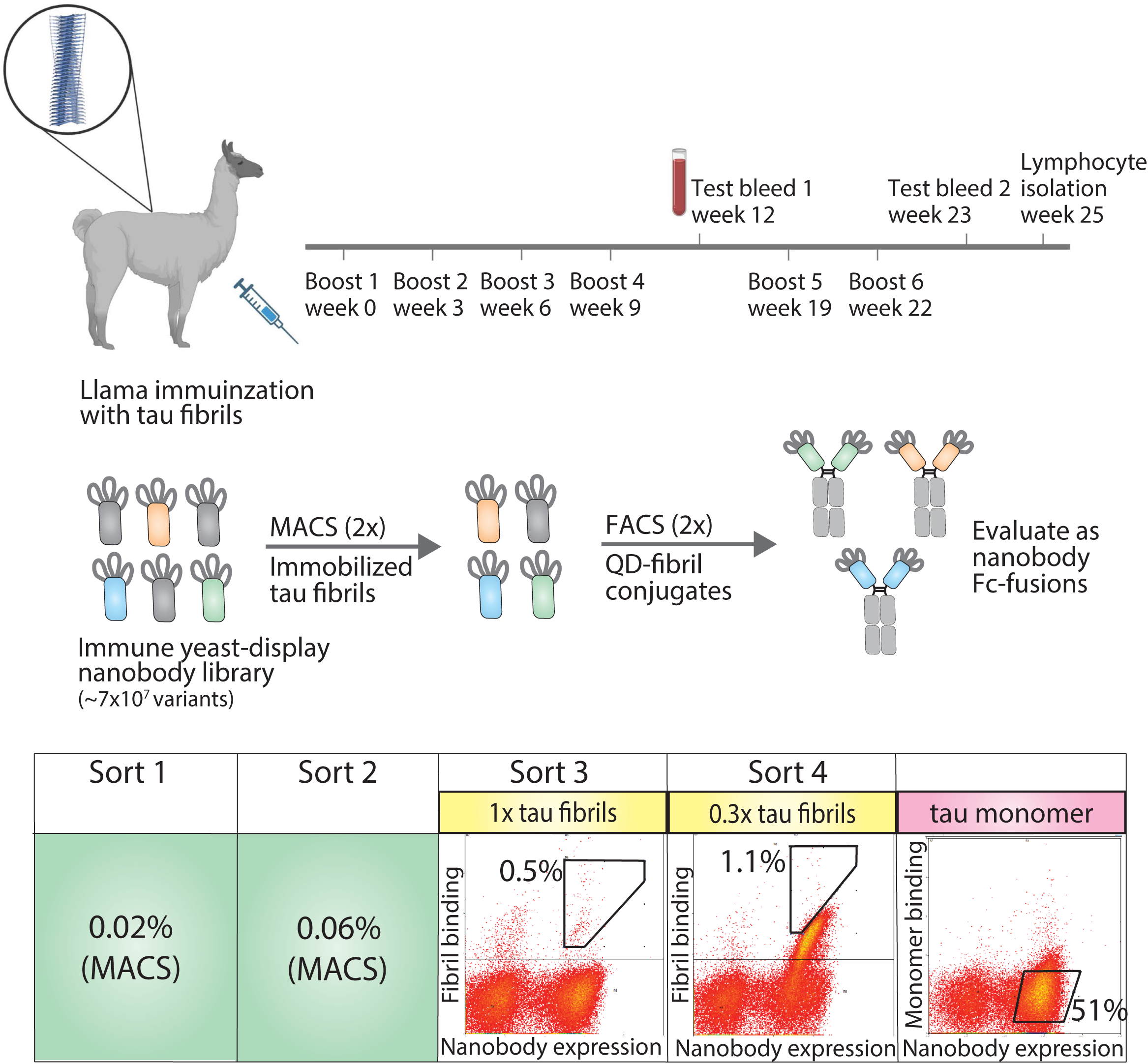
Overview of approach for isolating tau conformational nanobodies. A yeast surface display library was _rst prepared from a nanobody repertoire isolated after immunizing a llama with tau _brils. The library was sorted twice against tau _brils via magnetic-activated cell sorting (MACS) to initially enrich the library. Fluorescence-activated cell sorting (FACS) was then used to select a population of yeast cells that bound to tau _brils in a manner proportional to nanobody expression. Next, the enriched library was profiled for binding to tau monomer to evaluate conformational specificity. Finally, the enriched library was sequenced and selected clones were expressed as nanobody Fc-fusion proteins for evaluation. Cells were collected from gates with percentages labeled in sorts 3 and 4. The gate and percentage included on monomer profiling serve as a reference to demonstrate that the majority of yeast cells displaying nanobodies on their surface do not show binding signal for tau monomer.

The nanobody library was first sorted twice against HT40 fibrils using magnetic-activated cell sorting (MACS), and modest enrichment in the percentage of cells collected was observed between the first (0.02%) and second (0.06%) sorts. The enriched library was then further sorted twice for binding to tau fibrils using fluorescence-activated cell sorting (FACS). In these sorts, tau fibrils were captured on the surface of fluorescent quantum dots (QD) using a sequence-specific tau antibody (Tau-5) (32). Yeast cells that both displayed nanobodies (as detected using Myc-tag detection) and bound to antigen (as detected using QD fluorescence signal) were collected. In the third sort (FACS sort #1), a modest population of cells was collected that displayed antigen-binding signal (∼0.5%). In the fourth sort (FACS sort #2), strong enrichment for antigen-binding signal was observed, and a population of cells was collected that displayed antigen-binding signal in direct proportion to nanobody expression level. Finally, because we desired nanobodies that bind tau aggregates with conformational specificity, the binding of the enriched library to tau monomer was examined. The library displayed a minimal level of binding to tau monomer, and no further sorting was needed to reduce the level of tau monomer binding. Nanobodies were then Sanger sequenced from the fourth sort and selected for analysis. Three related nanobody sequences were observed, namely WA2.22, WA2.21, and WA2.7 (**Fig. S2**).

The three nanobodies were cloned as Fc-fusion proteins, expressed, and analyzed. They expressed at intermediate levels in HEK293-6E cells, with purification yields of 11-16 mg/L. The proteins displayed relatively high purity, as judged by both SDS-PAGE (**Fig. S3**) and size-exclusion chromatography (**Fig. S4**). Moreover, the affinities of the three selected nanobody Fc-fusion proteins were analyzed using a flow-cytometry based assay (33,34). Notably, all three bound tau aggregates (**Fig. 2A**), demonstrating that secondary screening was unnecessary to identify antigen-specific nanobodies. WA2.22 displayed the highest affinity of the three as a nanobody-Fc fusion protein (EC50 of 10.1±1.5 nM), which was approximately an order-of-magnitude higher than the two control mAbs (Tau-5 and zagotenemab; **Fig. S5**) generated using mouse immunization. The affinity of WA2.22 was also analyzed as a monovalent nanobody compared to its bivalent Fc-fusion counterpart, which revealed greater than an order-of-magnitude reduction in affinity as a monovalent nanobody (**Fig. S6**). This suggests that avidity is key to mediating the binding affinity of the bivalent nanobody Fc-fusion protein.

**Figure 2.**
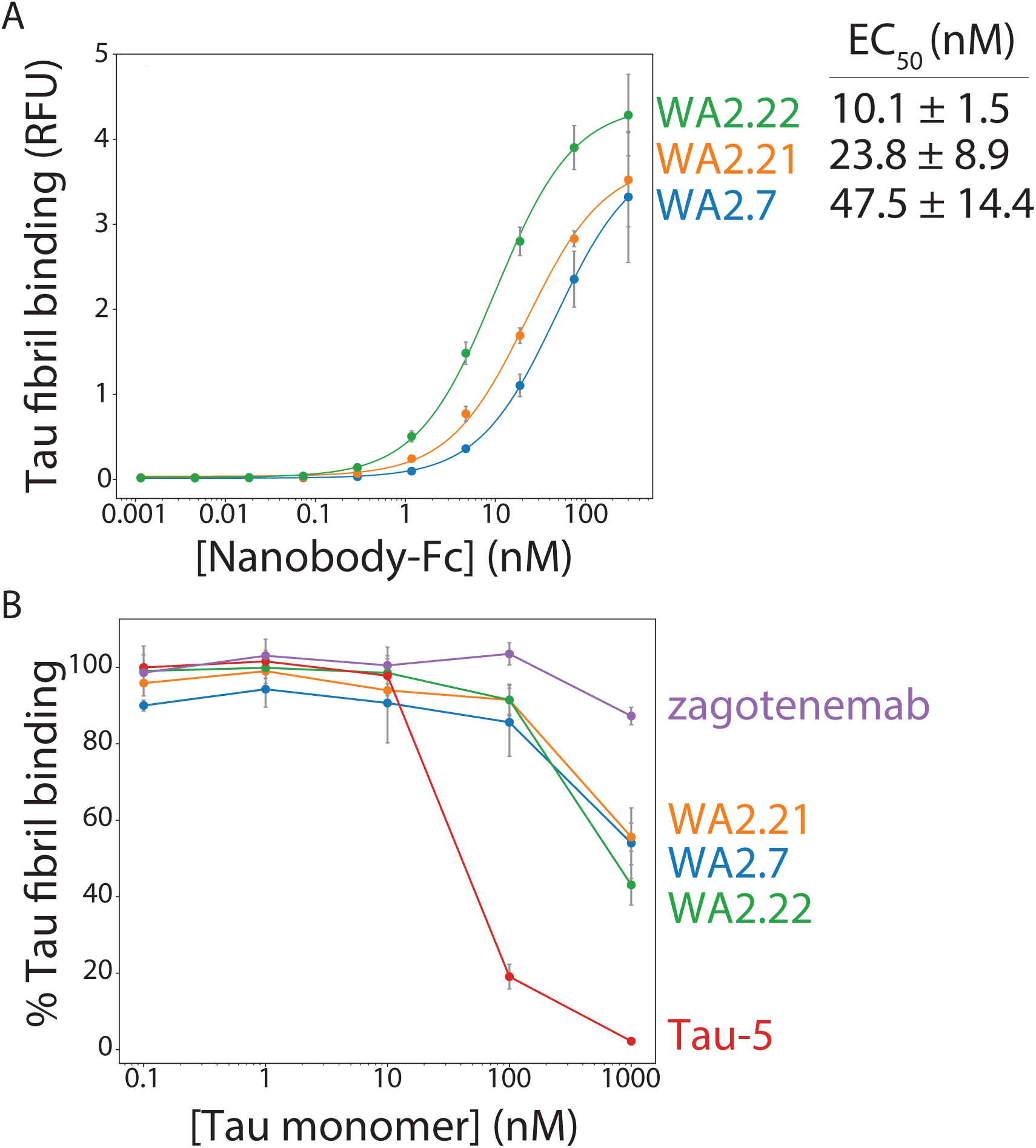
A_nity and conformational speci_city of selected tau nanobody Fc-fusion proteins. (A) Nanobody Fc-fusion proteins (WA2.22, WA2.21, and WA2.7) were incubated with tau _bril-coated magnetic beads at various concentrations. Nanobody binding was detected using an anti-human Fc 647 secondary antibody. Mean binding signal at each nanobody concentration was then determined using _ow cytometry. (B) Nanobody Fc-fusion proteins as well as two conventional antibodies (Tau-5 and zagotenemab), at a _xed concentration (10 nM), were _rst preincubated with tau monomer (0.1-1000 nM). Next, tau _bril-coated magnetic beads were then added to the mixture of antibody and tau monomer for approximately 3 h. Finally, nanobodies and antibodies bound to tau _bril-coated beads were detected via _ow cytometry, and the percentage of binding relative to that observed without tau monomer preincubation is reported. In (A) and (B) the data are averages, and the errors are standard deviations for three independent experiments…………………………………………………………………………………………………………………………….

The conformational specificity of the selected nanobodies was also examined, which was done by preincubating nanobody Fc-fusion proteins or control antibodies at a fixed concentration (10 nM) with various concentrations of tau monomer (0.1-1000 nM) before allowing them to bind immobilized tau fibrils. For comparison, a clinical-stage tau conformational antibody (zagotenemab) and a sequence-specific antibody (Tau-5) were included in this analysis. Tau-5 displays reduced binding to tau fibrils when the monomer concentration is in excess of the antibody concentration (**Fig. 2B**). At a 100-fold excess tau monomer concentration, Tau-5 retains only ∼2% of its binding to tau fibrils, and at a 10-fold excess monomer concentration, it retains only ∼19% of its binding. In contrast, a clinical-stage conformational antibody (zagotenemab) retains ∼87% of its binding at 100-fold excess tau monomer and maintains the entirety of its binding at all other monomer concentrations. Encouraging, the nanobody Fc-fusion proteins display conformational specificity for tau aggregates, as they retain 43-56% of their binding in the presence of 100-fold excess tau monomer and 86-91% of their binding in the presence of 10-fold excess tau monomer. These results demonstrate that the selected nanobodies recognize tau aggregates assembled from recombinant protein with conformational specificity.

### 3.2 Nanobody Fc-fusions recognize tau aggregates in mouse and human brain samples

After confirming the binding and conformational specificity of our selected nanobodies for recombinant tau fibrils, we next asked whether these nanobody Fc-fusion proteins selectively recognized tau aggregates formed *in vivo* in both a transgenic mouse model and human tauopathies. We began by analyzing their ability to recognize tau aggregates present in a transgenic P301S tau mouse model in comparison to wild-type (age-matched control) mice (**Fig. 3**). We evaluated the ability of two of our selected nanobody Fc-fusion proteins (WA2.21 and WA2.22), zagotenemab, and Tau-5 to recognize homogenized samples isolated from 11-month-old P301S transgenic or wild-type mice. As expected for a non-conformational antibody, Tau-5 binds to samples isolated from both P301S and wild-type mice. In contrast, our selected nanobodies and zagotenemab bind primarily to transgenic P301S samples.

**Figure 3.**
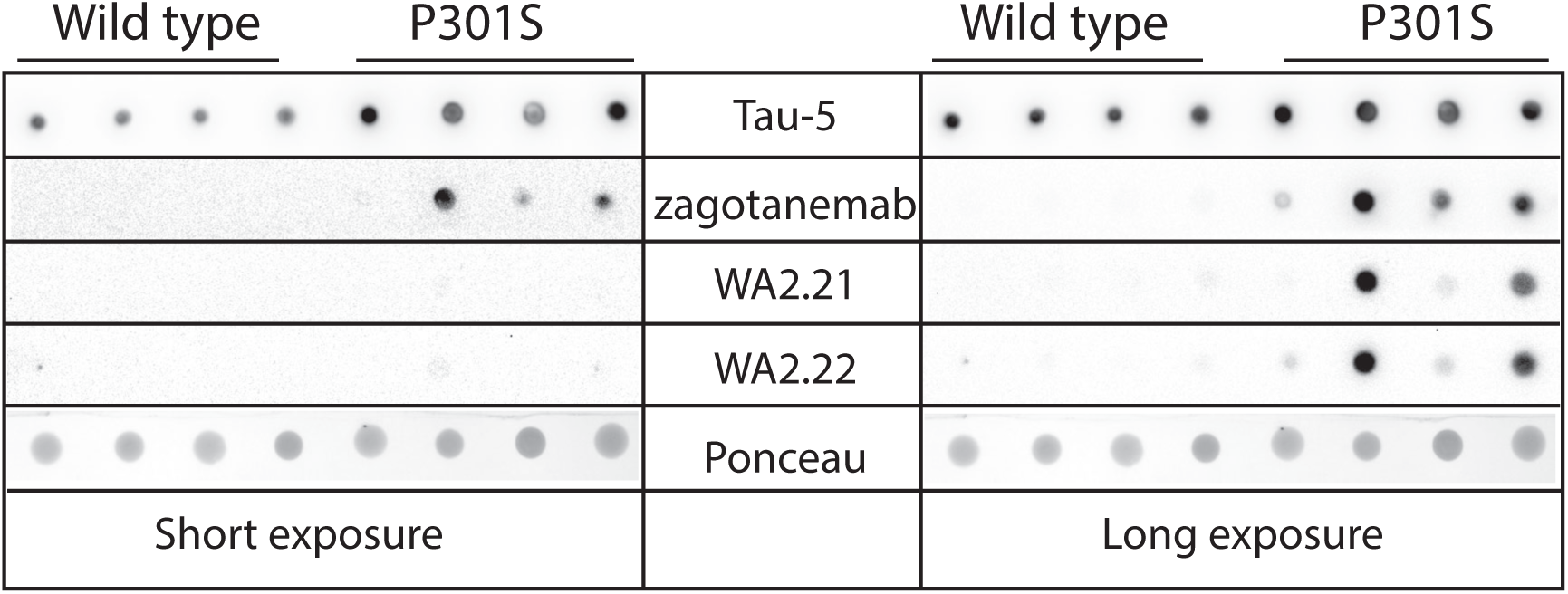
Immunodot analysis of tau conformational nanobodies using mouse brain samples. Immunodot blotting analysis of the selected nanobody Fc-fusion proteins (WA2.22 and WA2.21) was evaluated for both wild-type and transgenic P301S mouse brain homogenates. For comparison, a conformational tau antibody zagotenemab) and a sequence-specific tau antibody (Tau-5) were also analyzed. Immunoblots were imaged at both short (left) and long (right) exposure times. Ponceau stain was used as a loading control. The staining was repeated twice, and a representative image is shown.

Encouraged by these results, we next examined the ability of our highest affinity nanobody (WA2.22) to detect tau aggregates in mouse brain sections using immunostaining (**Fig. 4**). We stained both tissue sections from aged P301S transgenic mice and wild-type controls. For reference, we also stained these samples with a phospho-tau antibody (AT8) that recognizes tau aggregates in immunofluorescent staining (35). Importantly, we observed that WA2.22 Fc-fusion protein specifically stains transgenic tissue samples. Moreover, the WA2.22 staining co-localizes with AT8 staining, indicating that they stain similar tau aggregates in the transgenic mouse brain samples. Overall, our results indicate that WA2.22 displays conformational specificity for tau aggregates formed in the mouse brain.

**Figure 4.**
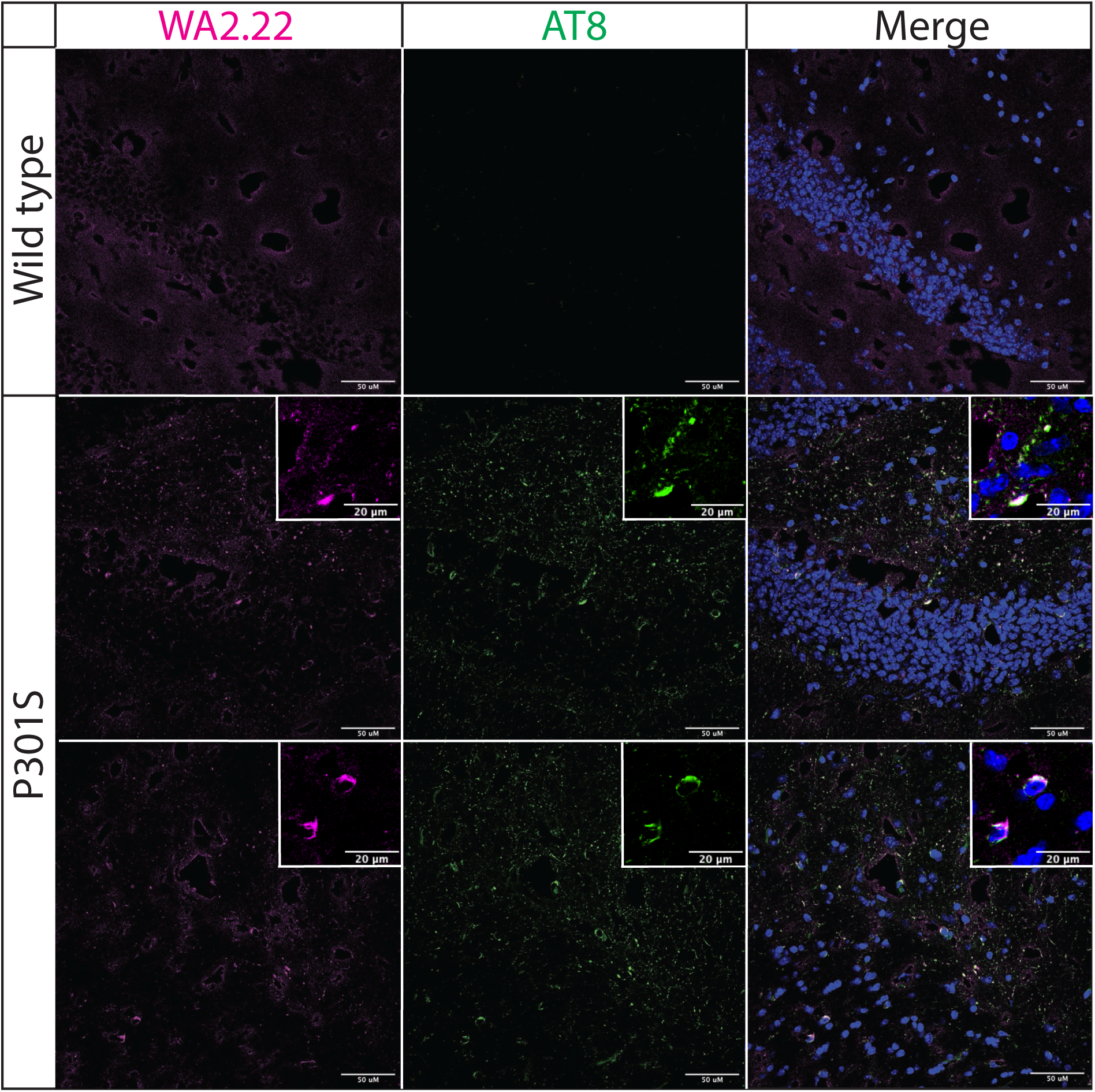
Immuno_uorescence analysis of a tau conformational nanobody using mouse brain samples. Immuno-_uorescent staining of _xed brain sections from wild-type and transgenic P301S mice was performed using WA2.22 purple; Fc-fusion protein). Tissue sections were co-stained with a phospho-tau antibody (AT8, green) and DAPI (blue). WA2.22 signal was detected using Alexa Fluor 647, and AT8 signal was detected using Alexa Fluor 488. The scale bars in the images represent approximately 50 μm, and the scale bars in the insets represent approximately 20 μm.

We also examined the ability of WA2.22 to stain tau aggregates in human tissue samples isolated from tauopathies in comparison to human tissue samples from subjects without cognitive impairment (**Fig. 5**). Encouragingly, we observed strong staining of WA2.22 Fc-fusion protein in tissue samples from both Alzheimer’s disease (AD) and progressive supranuclear palsy (PSP). Moreover, this staining strongly co-localized with the staining for AT8, and minimal signal was observed for either of these antibodies in control brains.

**Figure 5.**
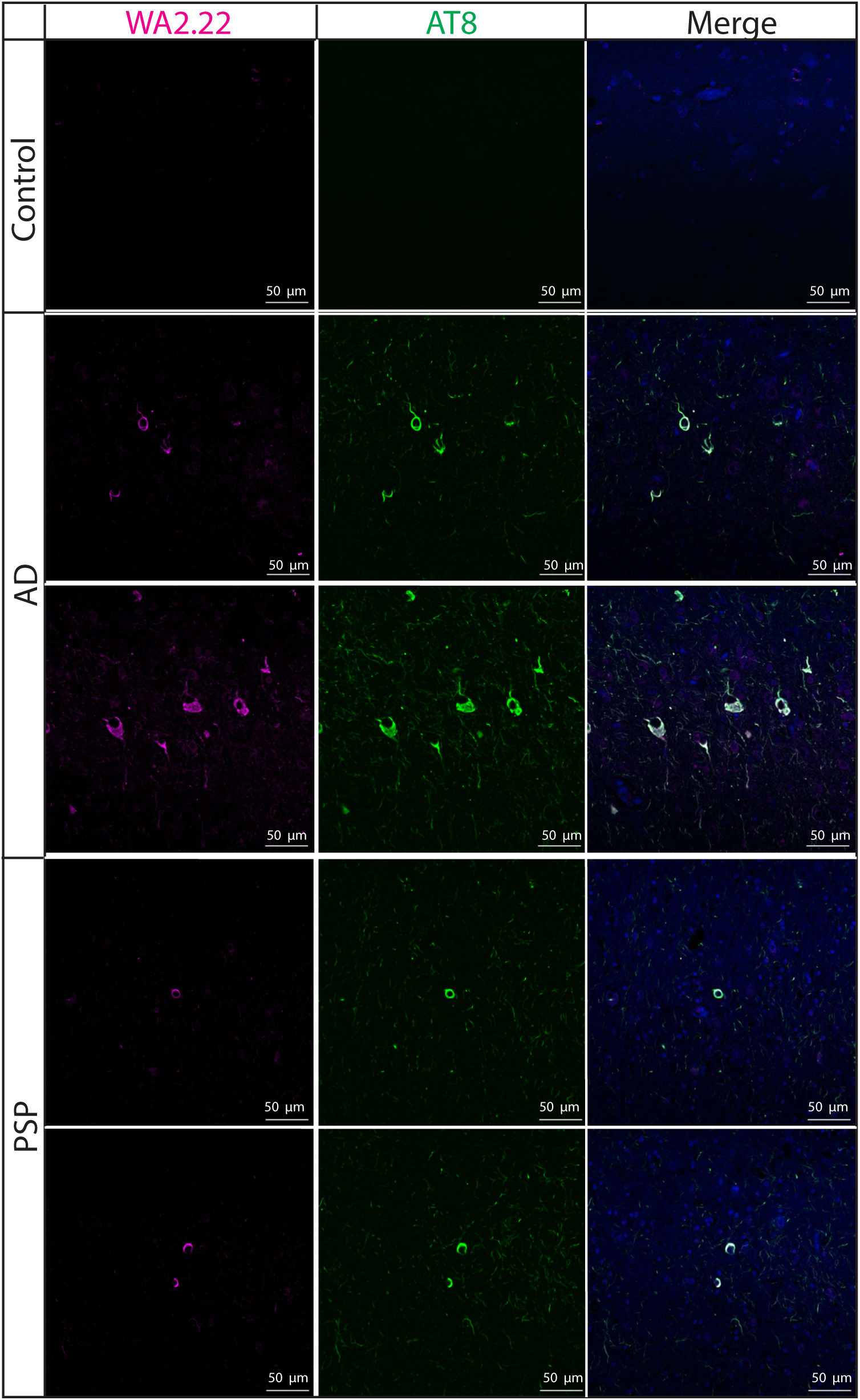
Immuno_uorescence analysis of a tau conformational nanobody using human brain samples. Immuno_uorescent staining of _xed brain sections from human samples without cognitive impairment (control), Alzheimer’s disease (AD), and progressive supranuclear palsy (PSP) was performed using WA2.22 (purple; Fc-fusion protein). Tissue sections were co-stained with a phospho-tau antibody (AT8, green) and DAPI (blue). WA2.22 signal was detected using Alexa Fluor 647, and AT8 signal was detected using Alexa Fluor 488. The scale bars in the images represent approximately 50 μm.

To complement the immunofluorescence staining, we also performed immunohistochemical staining of human brain tissue samples from Alzheimer’s disease using WA2.22 and zagotenemab (**Fig. 6**). We observed strong staining of tau aggregates by both WA2.22 Fc-fusion protein and zagotenemab. Further, we performed this analysis using adjacent brain sections for each of the two antibodies. We observe similar aggregate staining by both WA2.22 and zagotenemab in multiple locations throughout the analyzed brain sections. This result agrees with our observation of similar recognition of tau aggregates by WA2.22 and zagotenemab in mouse immunoblots (**Fig. 3**). Overall, our results demonstrate that WA2.22 shows strong conformational recognition of tau aggregates formed in human tauopathies by multiple methods.

**Figure 6.**
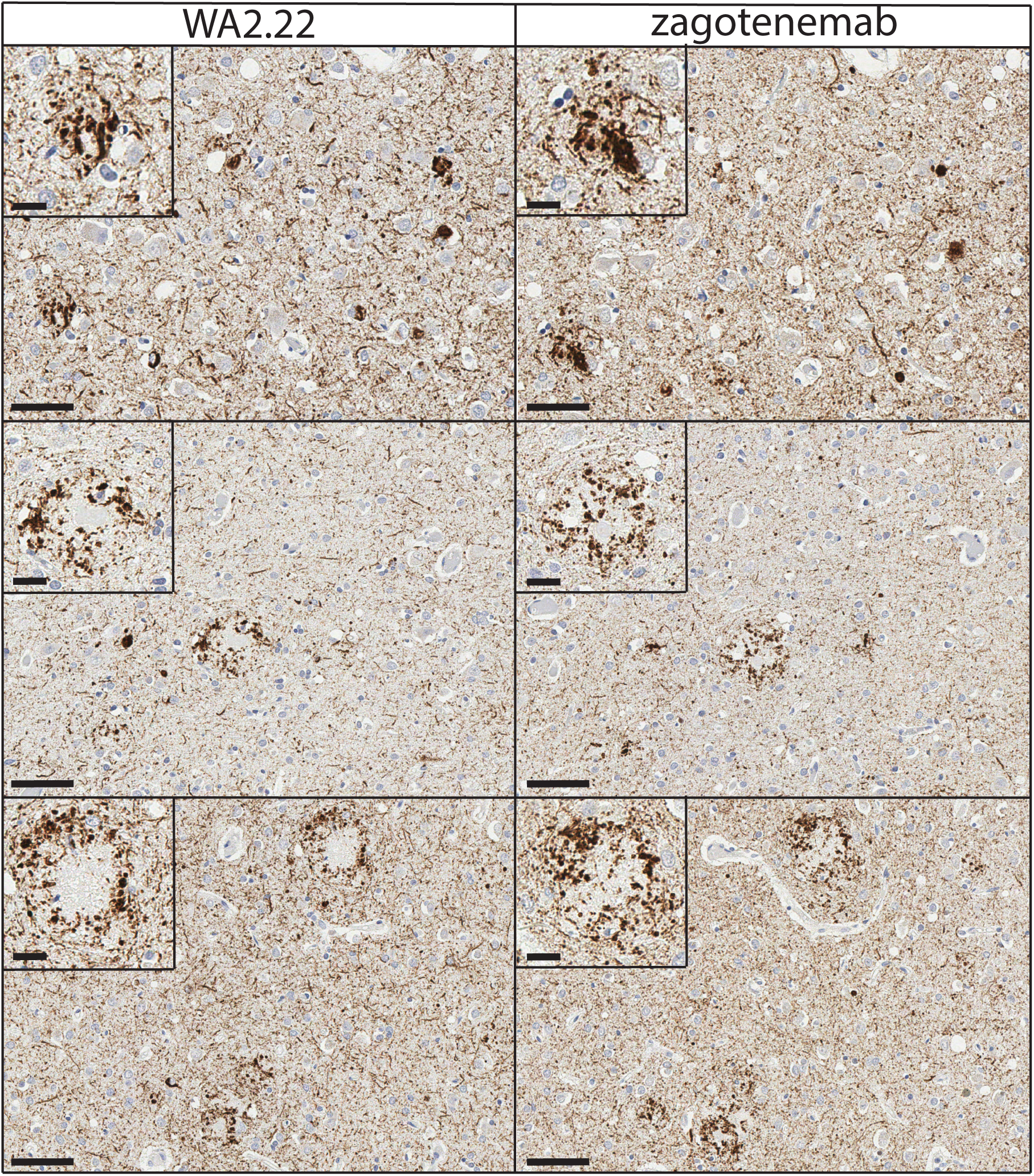
Immunohistochemistry analysis of a tau conformational nanobody using human brain samples. Immunohistochemical staining of _xed brain sections from human brain with a high level of Alzheimer’s disease neuropathological change (ADNC), NIA-AA criteria (A3, B3, C3) was performed using WA2.22 Fc-fusion protein (left) and zagotenemab (right). WA2.22 and zagotenemab staining was detected using horseradish peroxidase and developed with 3,3’-diaminobenzidine. Nuclei were detected via hematoxylin stain. The scale bars in the main images represent approximately 50 μm, and the scale bars in the insets represent approximately 20 μm.

### 3.3 Nanobodies display drug-like biophysical properties

To be highly useful in *in vivo* applications, such as diagnostic or therapeutic agents, nanobodies and antibodies more generally need to possess a combination of favorable biophysical properties, such as low non-specific binding (high specificity) and high stability, in addition to high affinity for their target antigen. Therefore, we examined the biophysical properties of our nanobody Fc-fusion proteins by first evaluating their non-specific binding to a polyspecificity reagent (**Figs. 7A** and **S7**). We used a polyspecificity reagent, namely soluble membrane proteins, prepared from the lysate of CHO cells (36). Interestingly, approved antibody drugs typically show lower levels of non-specific binding to this polyspecificity reagent than antibodies that are currently in clinical trials or that have failed in clinical trials (37). Notably, the tau nanobody Fc-fusion proteins demonstrate low non-specific binding and comparable levels to a highly specific clinical-stage antibody (elotuzumab). In contrast, zagotenemab, a conformational tau antibody originally generated via immunization (38), showed much higher non-specific binding than the nanobody Fc fusions and even higher levels than those for a clinical-stage antibody with previously reported high levels of non-specific binding (emibetuzumab) (37,39,40). For comparison, we also analyzed Tau-5, another antibody generated using immunization and found it also displays higher levels of non-specific binding than the nanobodies. These results indicate that our nanobodies show low non-specific binding in comparison to both clinical-stage controls and other tau antibodies.

**Figure 7.**
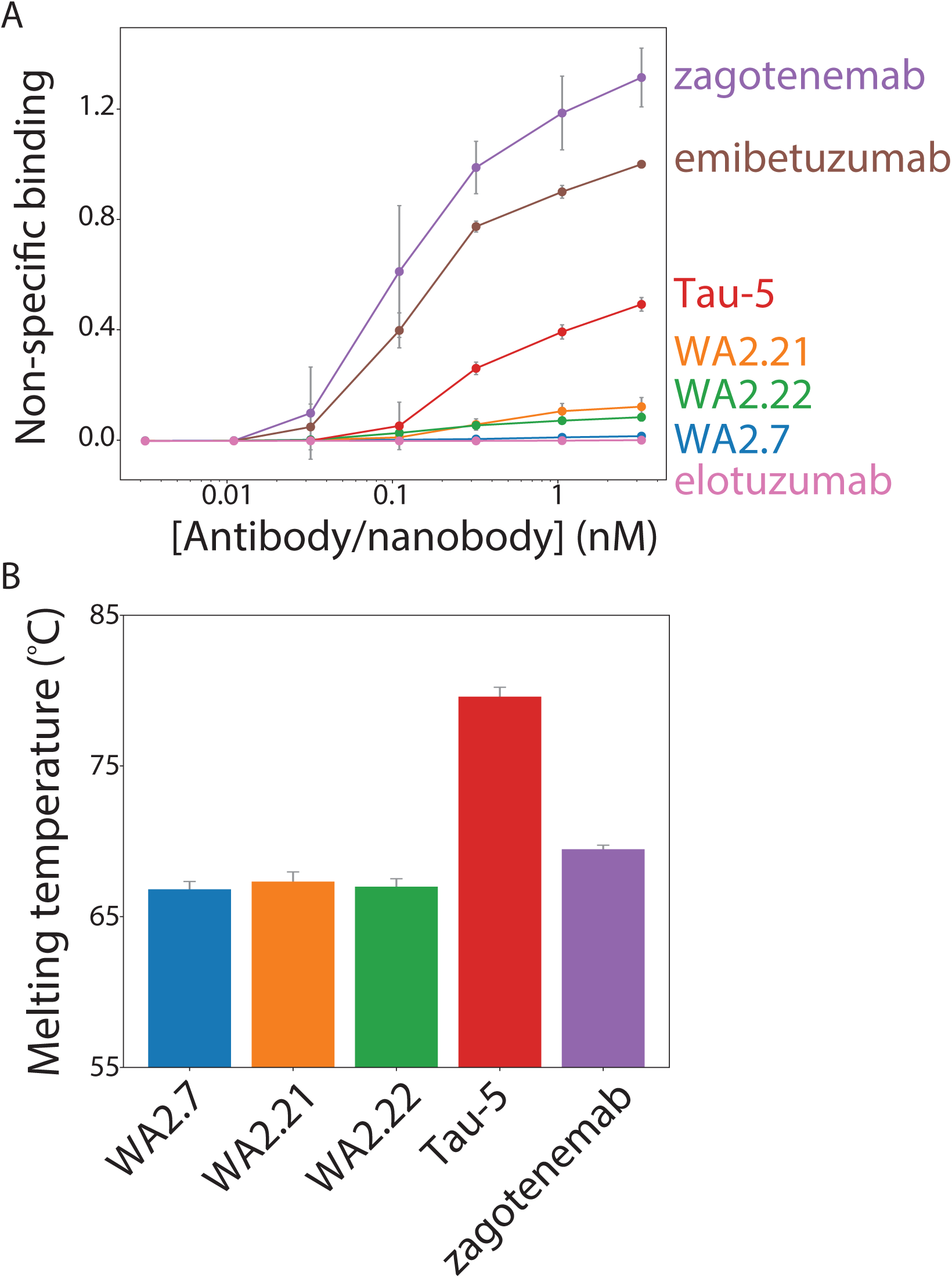
Biophysical characterization of tau conformational nanobodies. (A) Non-speci_c binding for the nanobod ies and antibodies was analyzed using a _ow cytometry assay. The nanobody Fc-fusion proteins and antibodies were immobilized on Protein A magnetic beads, and the levels of polyspecificity reagent binding (biotinylated soluble mem brane proteins from CHO cells) were evaluated using _ow cytometry. The measurements were normalized relative to two clinical-stage control antibodies with low (elotuzumab) and high (emibetuzumab) levels of non-specific binding. (B Nanobody Fc-fusion protein and antibody melting temperatures were analyzed by di_erential scanning _uorimetry. A single unfolding transition was observed, which is reported as the melting temperature. In (A) and (B), the data are aver ages, and the error bars are standard deviations for three independent experiments.

Finally, antibody stability is another key biophysical property of nanobodies and antibodies. Therefore, we analyzed the melting temperature (*Tm*) of the nanobody Fc-fusion proteins relative to conventional tau antibodies (**Fig. 7B**). Encouragingly, the nanobodies displayed melting temperatures of >65 °C (∼66.8-67.4 °C), which is a useful metric for identifying stable nanobodies (41,42). As expected, Tau-5 (*Tm* of 80.3 ± 0.6°C) and zagotenemab (*Tm* of 69.5 ± 0.3°C) showed higher stability due to the presence of stabilizing constant regions in these antibodies (CH1 and CL), which are absent in nanobody Fc-fusion proteins. Overall, these findings demonstrate that the tau conformational nanobodies in this work also have biophysical properties that are similar or better than those for clinical-stage antibodies.

## 4. DISCUSSION

We have demonstrated that tau conformational nanobodies can be readily isolated, without the need for any secondary screening, following llama immunization using a quantitative library sorting approach. The approach reported in this study has enabled the isolation of three nanobodies with related sequences (**Fig. S2**). These nanobodies share the same sequences of their CDR regions, but we observe overall differences in the binding properties and characteristics of the isolated nanobodies resulting from differences in their framework sequences. The majority of previous nanobodies generated via immunization have been selected using phage display (6,9,12,13,29,30), while few such immune libraries have been screened using yeast surface display (7,43,44). The application of yeast surface display to nanobody selection has been previously reported to result in a range of affinities for the isolated nanobodies depending on the library source (e.g., non-immune or immune) and sorting strategy (e.g., MAC-based or FACS-based), spanning sub-nanomolar affinities (44)to low nanomolar (43) to affinities >100 nM(7). Interestingly, the binding of the nanobodies in this study appears to be heavily influenced by the valency in which they are tested. While we have mainly examined the binding characteristics of nanobody Fc-fusion proteins in this study, testing of WA2.22 in a monovalent format indicates that the apparent affinity of this monovalent nanobody is greatly reduced compared to WA2.22 Fc-fusion protein (**Fig. S6**). This finding likely indicates that avidity, resulting from both the bivalency of the Fc fusion format and polyvalency of the aggregated tau antigen, plays a role in the interaction between these binding domains and tau aggregates.

Our unique methods for screening yeast-displayed libraries following immunization using FACS enables predictable isolation of nanobodies with a combination of desirable binding properties, including sequence and conformational specificity for tau aggregates. Our sorting process required only four total rounds of enrichment to directly isolate nanobodies with the desired properties. This report builds on our previous findings that QD immunoconjugates can be used to immobilize insoluble amyloid aggregates, which can then be used for library sorting in a similar manner as soluble antigens are used in conventional FACS sorting (32). Here, we further demonstrate the broad utility of this method and how it can be used for enriching an immune library in a surprisingly simple and predictable manner for directly isolating tau conformational nanobodies.

The nanobodies reported in this study should be considered in the context of similar antibodies and related nanobodies that have previously been reported. The vast majority of tau conformational antibodies reported up to this point have been conventional IgGs (45–49). These antibodies have been critical to studying differences in tau fibril morphology present in different tauopathies (45), understanding the progression of tau aggregation (46,47), and testing the effects of targeting tau aggregates in *in vivo* models of neurological disease (48–50). Similar to our findings, these antibodies have been reported to selectively recognize aggregates in mouse and human tissues (45–50). Our findings that our nanobody Fc-fusion proteins demonstrate conformational specificity for recombinant fibrils (**Fig. 2**), aggregates formed in P301S transgenic mouse tissue (**Figs. 3** and **4**), and aggregates present in Alzheimer’s disease (**Figs. 5** and **6**) and progressive supranuclear palsy (**Fig. 5**) tissue samples indicate that our nanobodies have potential for further evaluation and study of tau aggregates in neurodegenerative models.

More recently, nanobodies that target various forms of the tau protein have been reported in addition to conventional IgGs, including nanobodies targeting phospho-tau (28) and tau monomer (25–27). However, to the best of our knowledge, our tau nanobodies are the first reported conformational nanobodies that recognize tau aggregates. The only previously reported conformational nanobody specific for complex protein aggregates is one specific for α-synuclein (20), which is considerably smaller than tau (140 amino acids for α-synuclein versus 441 amino acids for the longest isoform of tau). The other conformational nanobodies reported previously typically recognize less complex peptide aggregates, such as those composed of Aβ (21,22,31).

Overall, the nanobody Fc-fusion proteins reported in this study have a combination of favorable binding and biophysical properties. It has previously been reported that nanobodies, and antibodies more generally, display trade-offs between interconnected properties, such as affinity, stability, and specificity (51). Encouragingly, the nanobodies generated in this study show a favorable combination of sequence specificity, conformational specificity, and high stability. It is particularly interesting that the nanobody Fc-fusion proteins demonstrate low non-specific binding relative to tau antibodies generated by traditional immunization methods. Tau-5 was isolated following mouse immunization, and zagotenemab is the humanized form of MC1, which was also generated via mouse immunization (38). While these antibodies have high affinity for tau (**Fig. S5**), they suffer from limitations in moderate to high off-target binding. This is also notable given that other well-known amyloid-specific antibodies that were evaluated in clinical trials, such as gantenerumab and aducanumab, also display high levels of non-specific binding (34,37), revealing that such antibodies have an increased risk for non-specific binding. In the future, it would be simple to incorporate negative flow cytometric selections for a lack of binding to polyspecificity reagents, which could be used for further ensure selection of conformational nanobodies and antibodies with low levels of non-specific binding.

## 5. CONCLUSIONS

We have reported tau conformational nanobodies with a combination of favorable binding and biophysical properties without the need for any secondary screening. The characteristics of the tau nanobodies suggest several potential future opportunities. First, while the nanobody Fc-fusion proteins reported here display affinities in the 10-50 nM range, it would be straightforward to further enhance their affinity using standard mutagenesis methods and our quantitative flow cytometric methods (42,52–55). Additionally, the ability of the tau conformational nanobodies to strongly and specifically recognize tau aggregates in mouse and human brain samples motivates their evaluation in biological assays or *in vivo* applications. Some advantages of evaluating nanobodies in such applications include their small size and modular nature, which has previously been reported to readily enable the incorporation of nanobodies into various multivalent and bispecific formats (41,56–59). Multivalent or bispecific nanobodies have many applications associated with neurodegenerative diseases. An attractive future direction would be to test these nanobodies in bispecific antibody shuttles that cross the blood-brain barrier to examine their antigen binding within the brain after intravenous administration. These and other potential applications of the conformational nanobodies, which we expect can be readily generated using the methods reported here, are expected to accelerate the study, detection, and potentially treatment of diverse neurodegenerative diseases.

## 6. MATERIALS AND METHODS

### 6.1 Llama immunization and immune library generation

The immunization protocol was performed under contract by Triple J Farms (Bellingham, WA) and was approved by Triple J Farms Institutional Animal Care and Use Committee (IACUC). An adult male llama named Walkabout was immunized with dGAE fibrils (StressMarq Biosciences, SPR-461). Walkabout received four injections of 200 μg of sonicated dGAE fibrils at 3-week intervals. A serum sample was collected following the fourth injection, and the presence of antibodies which bind to immobilized HT40 and dGAE fibrils and monomer analyzed by flow cytometry. Briefly, DynaBeads M-280 tosylactivated (Fisher, 14203) conjugated with fibrils, monomer, or unconjugated (background) were blocked with 10 mM glycine for 1 h and then washed once with 1x PBS plus 0.1% BSA (PBSB). The beads were then incubated with 10-fold dilutions of serum collected either before the first injection (pre-bleed) or after the fourth boost (test bleed 1). The incubation was performed at room temperature for approximately 3 h with mild agitation. Following the serum incubation, the beads were washed once with ice cold PBSB and incubated with a 1:300 dilution of goat anti-alpaca IgG H+L (also reactive for llama antibodies) Alexa Fluor 647 (Jackson ImmunoResearch, 128-605-160) on ice for 4 min. The beads were then washed once with ice cold PBSB, resuspended in PBSB, and analyzed by flow cytometry using a BioRad ZE5. The mean fluorescence signals were recorded, and values reported are normalized to the mean signal obtained from corresponding beads incubated without serum but with secondary antibody incubation. Following initial serum analysis, two additional boosts of 200 μg of sonicated dGAE fibrils were performed at 3-week intervals, serum was collected, and the presence of antibodies which bind to HT40 and dGAE fibrils and monomer was analyzed by flow cytometry in the same manner as previously described (pre-bleed and test bleed 2). Blood was collected and bulk lymphocytes were isolated by gradient centrifugation using Lymphoprep (Fisher, NC0418243). Lymphocytes were then frozen and stored at -80 °C for future use.

Lymphocytes were thawed, and RNA was extracted using a Macherey-Nagel NucleoSpin RNA kit (Fisher, NC9581114) according to the manufacturer’s protocol. Reverse transcription was then performed using Superscript III reverse transcriptase (Fisher, 18-080-044) and random primers (Fisher, 10-777-019) to generate cDNA. A first PCR was then performed using primers which anneal to the antibody leader sequence and CH2 domain (60). The PCR product was purified using a 2% agarose (Fisher, BP160-500) gel, and VHH sequences (band corresponding to ∼ 600 bp) were separated from VH sequences (band corresponding to ∼900 bp). VHH DNA was further amplified using primers that bind FR1 and FR4 or the long and short hinge of heavy-chain antibodies (61–63). A final PCR was performed to introduce overlap with a modified version of the pCTCON2 yeast-surface display plasmid for homologous recombination. A yeast-surface display library was prepared as previously described (33,54,64). Approximately 7.2 x 10^7^ transformants were obtained.

### 6.2 Material preparation

HT40 beads were prepared by sonicating 100 μg HT40 fibrils (StressMarq Biosciences, SPR-329) for 5 min (30 s on, 30 s off) in 500 μL of 20 mM HEPES. 8 x 10^7^ DynaBeads M-280 tosylactivated (Fisher, 14203) were washed twice with 1 mL of 20 mM HEPES. Washed beads were then mixed with 100 μg sonicated HT40 fibrils and allowed to incubate with end-over-end mixing for 2-3 d in a total volume of 1 mL 20 mM HEPES. Beads were stored at 4 °C until use.

dGAE beads were prepared by sonicating 100 μg dGAE fibrils (StressMarq Biosciences, SPR-461) for 5-10 min (30 s on, 30 s off) in 500 μL of 20 mM HEPES. 8 x 10^7^ DynaBeads M-280 tosylactivated (Fisher, 14203) were washed twice with 1 mL of 20 mM HEPES. Washed beads were then mixed with 100 μg sonicated dGAE fibrils and allowed to incubate with end-over-end mixing for 2-3 d in a total volume of 1 mL 20 mM HEPES. Beads were stored at 4 °C until use.

Quantum dot (QD)-capture antibody conjugates were prepared as previously described (32). A Site-click Qdot 655 antibody labeling kit (Invitrogen, S10453) was used to conjugate 125 μg of Tau-5 to dibenzocyclooctyne (DIBO) modified QDs. Conjugation was performed according to the manufacturer’s protocol, and QD-Tau-5 conjugates were stored at 4 °C until use.

### 6.3 Library sorting to identify tau nanobodies

Yeast cells displaying nanobodies were first enriched for nanobodies which bind to HT40 (full-length tau) fibrils using two rounds of MACS. In the first MACS selection, 1 x 10^9^ yeast cells were washed twice with PBSB. 1×10^7^ HT40 fibril-coated tosyl beads were blocked twice with 10 mM glycine and washed once with PBSB. Yeast cells were then mixed with prepared HT40 fibril-coated beads in a total volume of 5 mL PBSB with 1% milk. Yeast cells were incubated with HT40 fibril-coated beads for ∼3 h at room temperature with end-over-end mixing. Following this incubation, mixture was placed on a DynaMag-15 magnet (Invitrogen, 12301D), and beads and bound cells were washed once with 10 mL ice-cold PBSB. Yeast bound to HT40 fibril-coated beads were then transferred to a flask containing 50 mL SDCAA and allowed to grow at 30 °C for 2 d. Dilutions of the culture were plated immediately after performing MACS to estimate the number of cells collected. The second MACS selection was performed similarly except that 1 x 10^7^ yeast cells were used, and the final incubation volume was 1 mL.

The third and fourth sorts were performed using FACS as previously described (32). In sort 3, 5 μg of HT40 fibrils were sonicated for 5 min (30 s on, 30 s off), mixed with 5 μL QD-Tau5 conjugates, and incubated with end-over-end mixing for 2 h. 1 x 10^7^ yeast cells were washed twice with PBSB. Yeast cells were combined with QD-fibril complexes in a total volume of 200 μL with 1% milk and 1:1000 mouse anti-Myc antibody (Cell Signaling, 2276S) and allowed to incubate with end-over-end mixing at room temperature for approximately 3 h. Following this primary incubation, yeast cells were washed with ice-cold PBSB, incubated with 1:200 goat anti-mouse Alexa Fluor 488 (Invitrogen, A11001) and 1:1000 streptavidin Alexa Fluor (Invitrogen, S32357) on ice for 4 min, and washed with ice-cold PBSB. Immediately prior to sorting, cells were resuspended in ice-cold PBSB. Sorting was performed on a Beckman Coulter MoFlo Astrios sorter. Sort 4 was performed in the same manner as sort 3 except that QD-fibril complexes were prepared by sonicating 1.67 μg of HT40 fibrils and mixing with 1.67 μL QD-Tau5 conjugates.

Finally, the enriched library was examined for binding affinity toward HT40 monomer. 1×10^7^ yeast cells were washed twice with PBSB and incubated with 10 nM recombinant His-tagged HT40 monomer. Incubation was performed in a final volume of 1 mL with end-over-end mixing at room temperature for approximately 3 h. Following primary incubation, yeast cells were washed once with ice-cold PBSB. Yeast cells were incubated with 1:1000 dilution of mouse anti-Myc antibody and 1:1000 dilution of chicken anti-His (Invitrogen, PA1-9531) antibodies on ice for 20 min. The cells were then washed once with ice-cold PBSB, incubated on ice with a 1:200 dilution of goat anti-mouse AlexaFluor 488 and a 1:1000 dilution of donkey anti-chicken IgY F(ab)’2 Alexa Fluor 647 (Jackson ImmunoResearch, 703-606-155) on ice for 4 min, and washed once more with ice-cold PBSB.

### 6.4 Nanobody cloning and expression

Plasmids of enriched nanobodies were isolated from the terminal round of sorting using a yeast miniprep kit (Zymo, D2004). For nanobody Fc fusions, nanobody sequences were amplified by PCR, digested using NheI-HF (New England Biolabs, R3131L) and HindIII-HF (New England Biolabs, R3104S) restriction enzymes, and ligated (New England Biolabs, M0202L) into a nanobody Fc fusion (human IgG1 Fc) mammalian expression plasmid (pTT5). For monovalent WA2.22, the nanobody sequence was amplified by PCR to include a C-terminal 6x His-tag, digested using NheI-HF and BamHI (New England Biolabs, R3136S) restriction enzymes, and ligated into a mammalian expression plasmid (pTT5). Ligations were transformed into chemically competent DH5α *E. coli* cells. Cells were then plated on LB plates with ampicillin (100 μg/mL) selection marker and grown overnight at 37 °C. Individual colonies were then picked and grown in LB media supplemented with ampicillin (100 μg/mL) overnight at 37 °C. Plasmids were isolated using a bacterial miniprep kit (Qiagen, 27106). Nanobody sequences were determined by Sanger sequencing.

Nanobody Fc fusion proteins were expressed in HEK293-6E cells (National Research Council of Canada) via transient transfection. Monoclonal antibodies used in this study were all expressed with human IgG1 Fc and using the same expression and purification techniques as for the nanobody Fc-fusion proteins. Cell culture was carried using in F17 media (Invitrogen, A13835) supplemented with 0.1% kolliphor (Sigma-Aldrich, SLCL6020). Transfection was performed as previously described (65,66). Either 15 μg of nanobody Fc plasmid or 1.5 μg of nanobody Fc plasmid and 13.5 μg of ssDNA (Sigma, D7656) were combined with 45 μg PEI (Fisher Scientific, NC1038561) in 3 mL of F17 media, vortexed, allowed to incubate for 15 min, and added to cells. Approximately 24 h after transfection, protein expression was enhanced through the addition of 750 μL of 20% Yeastolate (Gibco, 292804). Cells were cultured for an additional 4-5 d and then harvested by centrifuging at 3500 xg for 40 min. For purification, approximately 300 μL of Protein A agarose beads (Thermo Scientific, 20334) was added to the supernatant and incubated overnight at 4 °C with mild agitation. The beads were recovered in a filter column (Fisher, 89898) and washed with 1x PBS. Proteins were eluted from Protein A beads by incubating with 0.1 M glycine (pH 3) and buffer exchanged into acetate buffer. Proteins were filtered, aliquoted, and stored at -80 °C until use.

Monovalent WA2.22 was expressed in HEK293-6E cells via transient transfection as described above. For purification, approximately 300 µL of Ni-NTA agarose beads (Qiagen, 30230) was added to the supernatant and NiSO4 was added to a final concentration of 1 mM. The supernatant was incubated with the beads over night at 4 °C with mild agitation. The beads were recovered in a filter column and washed with 1x PBS. The beads were then washed once with 50 mM imidazole (pH 7.4). WA2.22 nanobody was eluted from the beads by incubating with 500 mM imidazole (pH 7.4) and buffer exchanged into acetate buffer. The protein was filtered, aliquoted, and stored at -80 °C until use.

### 6.5 Antibody purity and analytical size-exclusion chromatography analysis

Nanobodies and antibodies were analyzed via size-exclusion chromatography (SEC) with a Shimadzu Prominence HPLC system. Following Protein A purification, nanobodies and antibodies were stored in 20 mM potassium acetate buffer (pH 5.0). Antibodies and nanobodies were diluted to 0.1-0.2 mg/mL in either 100 mM sodium acetate buffer (pH = 5.0) or 1x PBS (pH 7.4), and 100 μL was injected into a SEC column (Superdex 200 Increase 10/300 GL column; GE, 28990944). SEC analysis and purification was performed at 0.75 mL/min using a running buffer of either 100 mM sodium acetate and 200 mM arginine (pH 5.0) or 1x PBS and 200 mM arginine (pH 7.4). Absorbance was monitored at 280 nm, and the percentage of monomer was calculated using absorbance peaks between the void volume and buffer elution times. Nanobodies or antibodies which displayed below 90% monomer following Protein A purification were further purified by SEC, and proteins were further analyzed to ensure >90% monomer following SEC purification.

### 6.6 Nanobody Fc-fusion protein affinity analysis

Nanobody Fc-fusion protein affinity was analyzed using a bead-based flow cytometry assay (33,34). HT40 fibril-coated tosyl Dynabeads were blocked with 10 mM glycine with end-over-end mixing at room temperature for 1 h. Beads were then washed with PBSB. Immediately before use, nanobody Fc fusions were thawed at room temperature and centrifuged in a tabletop centrifuge at max speed (21,300 xg) for 5 min. The supernatant was transferred to a new tube, and the nanobody Fc fusion concentration was determined by measuring the A280 using a NanoDrop. Varying concentrations of nanobody Fc fusion (300 nM and 4x dilutions) were added to individual wells of a 96-well plate (Greiner, 650261) and incubated with 1×10^5^ prepared HT40 fibril beads in 1% milk. Incubation was performed for approximately 3 h at room temperature with mild agitation. Following primary incubation, the plate was centrifuged at 2500 xg for 5 min, and the beads were then washed once with ice cold PBSB. The beads were then incubated with a 1:300 dilution of goat anti-human Fc Alexa Fluor 647 (Jackson ImmunoResearch 109-605-098) in PBSB on ice for 4 min. The beads were then washed once with ice cold PBSB, resuspended in PBSB, and mean fluorescence signal was examined by flow cytometry using a BioRad ZE5 analyzer. The affinities of Tau-5 and zagotenemab were analyzed in the same manner.

### 6.6 Comparison of monovalent and bivalent WA2.22 affinity

The affinity of monovalent WA2.22 (6xHis tag at C-terminus) and bivalent WA2.22 Fc-fusion protein (6x His-tag and a FLAG-tag at C-terminus) was analyzed using a bead-based flow cytometry assay. HT40 fibril-coated tosyl Dynabeads were blocked with 10 mM glycine with end-over-end mixing at room temperature for 1 h. The beads were then washed once with PBSB. Immediately before use, WA2.22 nanobody and WA2.22 nanobody Fc-fusion protein were thawed at room temperature and transferred to a new tube, and the nanobody or nanobody Fc fusion concentration was determined by measuring the A280 using a NanoDrop. Varying concentration of monovalent WA2.22 (1000 nM and 4 x dilutions) and WA2.22 Fc fusion (250 nM and 4x dilutions) were added to individual wells of a 96-well plate and incubated with 1×10^5^ prepared HT40 fibril beads in 1% milk. Incubation was performed for approximately 3 h at room temperature with mild agitation. Following incubation with monovalent WA2.22 and bivalent WA2.22 Fc fusion, the plate was centrifuged at 2500 xg for 5 min, and the beads were washed once with ice cold PBSB. The beads were then incubated with a 1:1000 dilution of chicken anti-His antibody (Invitrogen, PA1-9531) on ice for 20 min. The beads were then washed once with ice cold PBSB. The beads were then incubated with a 1:1000 dilution of donkey anti-chicken IgY F(ab)’2 Alexa Fluor 647 (Jackson ImmunoResearch, 703-606-155) on ice for 4 min. The beads were then washed once more with ice cold PBSB, resuspended in PBSB, and mean fluorescence signal was examined by flow cytometry using a BioRad ZE5 analyzer.

### 6.7 Nanobody conformational specificity analysis

The conformational specificity of nanobody Fc-fusion proteins was analyzed using a bead-based flow cytometry assay (33,34). For comparison, a sequence specific antibody (Tau-5) and a highly conformational antibody (zagotenemab) were included in analysis. Nanobody Fc fusions or antibodies at a fixed concentration (10 nM) were first incubated with HT40 monomer at varying concentrations (0.1-1000 nM) in individual wells of a flow plate. Nanobody Fc fusion or antibody was also incubated under the same condition without monomer for comparison. Incubation was carried out in PBSB plus 1% milk for approximately 1 h at room temperature with mild agitation. HT40 fibril-coated beads were blocked and washed as described above, and 1×10^5^ beads were added to each well. After adding beads, incubation was performed for approximately 3 h at room temperature with mild agitation. Following incubation, the plate was centrifuged at 2,500 xg for 5 min, and the beads were washed once with PBSB. The beads were then incubated with a 1:300 dilution of goat anti-human Fc Alexa Fluor 647 (Jackson ImmunoResearch, 109-605-098) in PBSB on ice for 4 min. The beads were then washed once with ice cold PBSB, resuspended in PBSB, and mean fluorescence signal was examined by flow cytometry using a BioRad ZE5 analyzer. Percent binding was determined by comparing the mean fluorescence signal at a given monomer concentration to mean fluorescence signal in the absence of monomer.

### 6.8 Immunoblotting of mouse brain samples

All experiments were approved by the University of Michigan IACUC and performed in accordance with the National Institutes of Health Guide for the Care and Use of Laboratory Animals. The facility in which experiments were conducted was approved by the American Association for the Accreditation of Laboratory Animal Care. Mice were housed at the University of Michigan animal care facility. Mice were maintained according to a 12 h light/dark cycle with food and water available ad libitum (U.S. Department of Agriculture standard). Two strains of mice were bred at the University of Michigan: Hemizygous P301S tau mice (B6;C3-Tg-Prnp-MAPT-P301S PS19Vle/J; The Jackson laboratory stock #008169) (67) and non-transgenic littermates. Mice were euthanized at 9 and 11 months for sample collection.

For immunodot blotting, mouse brain homogenates were prepared as follows. Brain tissue from both 11-month-old P301S transgenic mice and age-match wildtype mice were first diluted in PBS at a 1:3 tissue:PBS ratio (w/v). Tissue in PBS was supplemented with a protease inhibitor cocktail and homogenized (Sigma Aldrich, 11873580001). Homogenized tissue was next centrifuged at 4 °C at 9,300 xg for 10 min. The supernatant was removed, and the pellet was resuspended in PBS with a second protease inhibitor cocktail (Roche, 11836170001). The resuspended pellet was then again centrifuged at 4 °C at 9,300 xg for 10 min. Following centrifugation, the supernatant was again removed, and the pellet was resuspended in 1% sarkosyl with protease inhibitor. The resulting mixture was vortexed for 1 min and then incubated at room temperature for 1 h. The mixture was then sonicated for 5 min using a water bath sonicator. These samples were then centrifuged at 4 °C at 16,000 xg for 30 min. From these samples, sarkosyl insoluble fractions of brain extract (7 µg of total protein) were spotted (1 µL) directly onto 0.45 µm nitrocellulose membranes and allowed to dry for 1 h. Loading controls were then stained with Ponceau S for 10 min and washed three times with distilled water. Membranes used for the analysis of tau nanobody Fc fusions and antibodies were blocked with 10% nonfat dry milk in Tris buffered saline supplemented with 0.1% Tween-20 (TBST) buffer.

Immunoblots were next incubated with nanobody Fc fusion proteins or antibodies at 10 nM. Incubation was carried out overnight at 4 °C in 1% milk in TBST. The immunoblots were next washed for 10 min, three times each with TBST. Immunoblots were next incubated with a HRP-conjugated goat anti-human IgG (1:5000 dilution) at room temperature for 1 h. Following this secondary incubation, the immunoblots were again washed three times, 10 min each with TBST. Immunoblots were then developed with an EcoBright Pico HRP Substrate (Innovative Solutions). Imaging was performed with a Genesvs G:Box imaging system (Syngene). Two independent repeats were performed.

### 6.9 Immunofluorescent staining of mouse brain samples

Brain tissue sections from 9-month-old P301S mice and age-matched non-transgenic controls were post fixed in methanol for 10 min, washed three times for 10 min each in 1x PBS, and subjected to heat-induced antigen-retrieval in 10 mM citrate buffer (pH 6) for 4 min. Brain sections were then washed twice with 1x PBS. Next, the brain sections were permeabilized by incubating for 10 min in 0.5% Triton-X 100. Following permeabilization, the sections were washed once with 1x PBS for 10 min. The brain sections were then blocked for 1 h using a Mouse on Mouse (M. O. M.) Blocking Regent (M.O.M. Immunodetection Kit, Vector, BMK-2202). After blocking, the brain sections were washed twice with 1x PBS for 2 min each. The sections were then incubated with M. O. M. diluent for 5 min. Next, the brain sections were incubated with both WA2.22 Fc fusion (100 nM) and AT8 (1:200 dilution, Invitrogen) in M. O. M. diluent at 4 °C overnight. The following day, the brain sections were washed three times with 1x PBS for 10 min each. Following washing, the brain sections were incubated for 1 h with goat anti-human IgG Alexa Fluor 647 (Invitrogen) and goat anti-mouse IgG Alexa Fluor 488 (1:500, Invitrogen). The brain sections were then washed three times with 1x PBS for 10 min each. The sections were then incubated with DAPI (Sigma) for 5 min at room temperature. The brain sections were then washed three times with 1x PBS for 5 min each. Finally, the brain sections were mounted with Prolong Gold Antifade Reagent (Invitrogen). Slides were imaged using an Olympus FV3000.

### 6.10 Immunofluorescent staining of human brain samples

Paraffin-embedded brain tissue sections from the frontal cortex of subjects with Alzheimer’s disease and progressive supranuclear palsy as well as age and gender matched controls were obtained from the Michigan Brain Bank (University of Michigan, Ann Arbor, MI, USA). Autopsy consent had been obtained from persons evaluated in the clinic and/or research studies; upon death of the individual, consent to autopsy was confirmed by next of kin. Samples were examined at autopsy, and neuropathological diagnosis was determined by University of Michigan Pathology Department neuropathologists. All protocols were approved by the University of Michigan Institutional Review Board and follow the declaration of Helsinki principles.

Brain sections were first heated, deparaffinized, and rehydrated through sequential washes with dilutions of xylene, ethanol, and distilled water. The brain sections were then subjected to microwave heat-induced antigen-retrieval in 10 mM citrate buffer (pH 6) for 4 min. Following antigen retrieval, the brain sections were permeabilized with 0.5% Triton-X 100, washed with 70% ethanol for 5 min, and then incubated with an autofluorescence eliminator reagent (Millipore catalog #2160) for 5 min. Next, the brain sections were washed three times with 70% ethanol. The brain sections were then blocked with a solution of 5% goat serum in 1x PBS for 1 h. The sections were then incubated with WA2.22 Fc fusion (100 nM) and AT8 (1:200, Invitrogen) overnight at 4 °C. On the following day, the sections were washed three times with 1x PBS for 10 min each. The brain sections were then incubated with goat anti-mouse Alexa Fluor 488 and goat anti-human Alexa Fluor 647 (1:500, Invitrogen). Following secondary staining, the sections were then washed three times with 1x PBS for 10 min each. The brain sections were then incubated with DAPI (Sigma) at room temperature for 5 min. Finally, the sections were washed three times with 1x PBS for 5 min each and mounted with Prolong Gold Antifade Reagent (Invitrogen). Slides were imaged using a Nikon A1 High Sensitivity Confocal (housed in the University of Michigan Biomedical Research Core Facilities Microscopy Core).

### 6.11 Immunohistochemical staining of human brain samples

Paraffin-embedded brain tissue sections from the frontal cortex of human brain with a high level of with Alzheimer’s disease neuropathological change (ADNC) NIA-AA criteria (A3, B3, C3) (68), were obtained from the Michigan Brain Bank as described above. Immunohistochemical staining was performed in the University of Michigan Rogel Cancer Center Histology core on the DAKO Autostainer Link 48 (Agilent, Carpiteria, CA). Tissue staining was performed at room temperature using a Human-on-Human HRP-Polymer kit (Biocare Medical, BRR4056KG). Briefly, WA2.22 Fc fusion and zagotenemab both with human IgG1 Fc were tagged with Digoxigenin for detection. Brain sections were deparaffinized in xylene, rehydrated through graded alcohols to water, and rinsed in TBS. Heat induced epitope retrieval was performed using Dako Envision Flex TRS,. Low pH peroxidase block was then applied to the slides for 5 min.. Digoxigenin-tagged WA2.22 Fc fusion or zagotenemab was then applied to slides and incubated for 60 min. Slides were rinsed with TBS and incubated with mouse anti-Digoxigenin secondary antibody for 15 min. Slides were rinsed with TBS and incubated with MACH 2 mouse HRP-polymer for 30 min. The slides were then rinsed with TBS and incubated with 3,3’-diaminobenzidine (DAB) for 10 min. Slides were rinsed with DI water, counterstained with hematoxylin, washed with DI water, and dehydrated through graded alcohols. Slides were cleared in xylene and coverslipped. The Digital Pathology slide scanning service, part of the Department of Pathology, Michigan Medicine, provided assistance with generation of whole-slide images.

### 6.12 Polyspecificity analysis

Biotinylated soluble membrane protein (SMP) reagent was prepared from CHO cells for polyspecificity analysis as previously described (36,69) and stored at -80 °C until use. Antibodies and nanobody Fc fusions were analyzed at equivalent molar concentrations across a range of concentrations. The assay was performed as previously described (69). The data from three independent repeats were normalized according to control antibodies with high (emibetuzumab) and low (elotuzumab) non-specific binding at the highest antibody or nanobody-Fc fusion concentration evaluated. Normalization is performed by setting the value of non-specific binding at the highest antibody concentrations to 1 for emibetuzumab (high non-specific binding) and 0 for elotuzumab (low non-specific binding), and scaling all other values accordingly.

### 6.13 Nanobody-Fc fusion melting temperature analysis

Nanobody-Fc fusion and antibody melting temperatures were analyzed using differential scanning fluorimetry. Nanobody-Fc fusion proteins and antibodies were diluted to a concentration of 0.12 mg/mL, and Protein Thermal Shift Dye (Applied Biosystems, 4461146) was to achieve a final concentration of 1x dye. A total of 20 μL protein and dye mixture was added to individual wells of a 394-well plate. Plates were submitted to the University of Michigan Advanced Genomics Core for analysis. A temperature gradient between 25 °C and 98 °C was examined. Three independent repeats were analyzed using a QuantStudio Real-Time PCR System.

### 7. LIST OF ABBREVIATIONS

QD: quantum dot
FACS: fluorescence-activated cell sorting
MACS: magnetic-activated cell sorting
CDR: complementarity-determining region
Tm: melting temperature
Fc: fragment crystallizable
BSA: bovine serum albumin
PBS: phosphate buffered saline
PBSB: PBS supplemented with 1% BSA
SEC: size-exclusion chromatography

## 8. DECLARATIONS

**Ethics approval and consent to participate.** Human brain tissue was collected with patient consent and protocols were approved by the Institutional Review Board of the University of Michigan and abide by the Declaration of Helsinki principles. Mouse brain samples were collected from mice with approval by the University of Michigan IACUC and performed in accordance with the National Institutes of Health Guide for the Care and Use of Laboratory Animals. The facility in which experiments were conducted was approved by the American Association for the Accreditation of Laboratory Animal Care. Finally, the llama immunization protocol was performed under contract by Triple J Farms (Bellingham, WA) and was approved by Triple J Farms Institutional Animal Care and Use Committee (IACUC).

**Consent for publication.** Not applicable.

**Availability of data and materials.** The datasets used and/or analyzed during the current study are available from the corresponding author on reasonable request.

**Competing interests.** None.

**Funding.** This work was supported by the National Institutes of Health (RF1AG059723 to P.M.T and R.S.K and R35GM136300 to P.M.T.; T32 NS007222 and F32 AG079576 fellowships to M.J.L; NIH P30 CA046592 to the University of Michigan Research Histology and Immunohistochemistry Core) and National Science Foundation (CBET 1159943, 1605266 and 1813963 to P.M.T., Graduate Research Fellowship to M.D.S.), the Albert M. Mattocks Chair (to P.M.T).

**Authors’ contributions.** J.M.Z. and P.M.T. designed the research, J.M.Z generated the immune library and QD-amyloid conjugates and performed the antibody library sorting, J.M.Z. and M.J.L. produced and/or purified the antibodies, J.M.Z, H.T., M.D.S., M.E.S., E.K.M. characterized the antibodies, H.T, M.E.S., M.D.S., J.M.Z., S.P.F., and H.L.P performed and/or analyzed the immunostaining results, M.D.S analyzed antibody binding data, and J.M.Z. and P.M.T. wrote the paper with input from the co-authors. All authors read and approved the final manuscript.

## Acknowledgements

**Acknowledgements.** We thank members of the Tessier lab for their helpful suggestions. We thank Matthias Truttmann for assistance with the primers used for identifying the primers used in immune library generation. We thank Kathy Toy from the University of Michigan Histology core for assistance with IHC staining. We thank Peter Ouillette from the Digital Pathology slide scanning service for assistance with imaging of IHC slides. Parts of Fig.1 were created with BioRender.com.

