## Supplemental Figures for "Quantitative flow cytometric selection of tau conformational nanobodies specific for pathological aggregates"

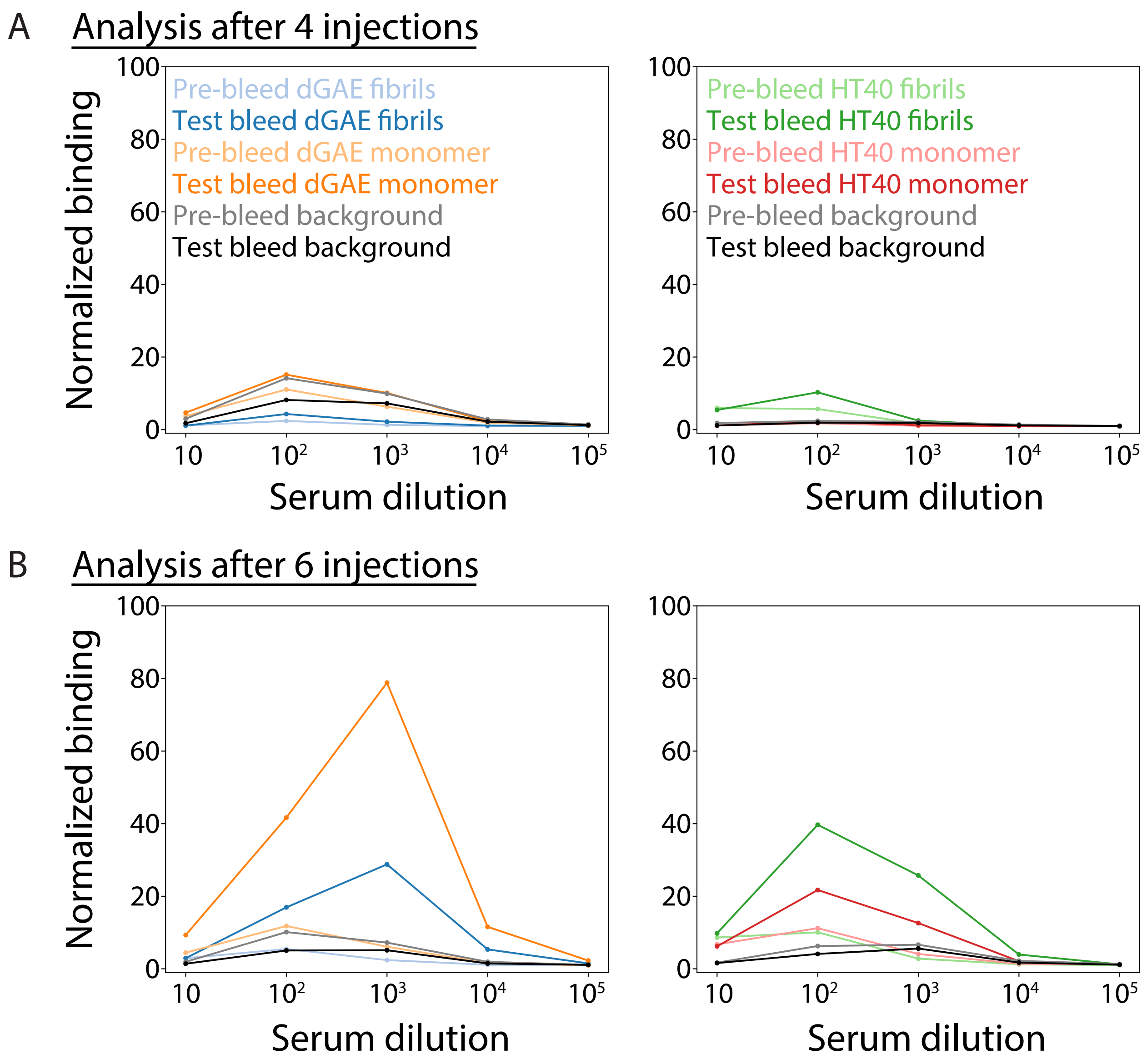


**Supplemental Figure 1. Analysis of llama serum for antibody binding to tau.** Llama serum was tested for the presence of antibodies that bound to HT40 and dGAE tau fibrils and monomers after (A) four and (B) six boosts of 200 µg of dGAE fibrils. Signal from various dilutions of serum was tested from test bleeds collected at multiple time points compared to the signal observed before the first injection (pre-bleed). The binding of antibodies in serum to magnetic tosyl beads coated with tau fibrils and monomer as well as beads blocked with glycine (background) was detected using an anti-alpaca/anti-llama IgG secondary. Signals are reported normalized to the signal observed from secondary incubated with each respective preparation of beads in the absence of serum. Serum testing was performed once in order to inform timing of lymphocyte isolation.

```
                                                 CDR1
        1                        26        35
WA2.22  DVQLQASGGGLVQPGGSLRLSCAAS GRIVSLGNMA WYRQAPGKQ
WA2.21  E........................ .......... .........
WA2.7   E........................ .......... .........

                  CDR2
             50              65
WA2.22  RELVA SVGRGGNTYYADSVKG RSTISRDDAKKMVALEMNSLKPE
WA2.21  ..... ................ ...........V...........
WA2.7   ..... ................ .......................

                    CDR3
                 98       107       118
WA2.22  DTAVYNCFV LVVQPTYDPY WGQGTQVTVSS
WA2.21  .....Y... .......... F..........
WA2.7   ......... .......... F..........
```

**Supplemental Figure 2. Amino acid sequences of tau conformational nanobodies.** Nanobody sequences were determined by Sanger sequencing.

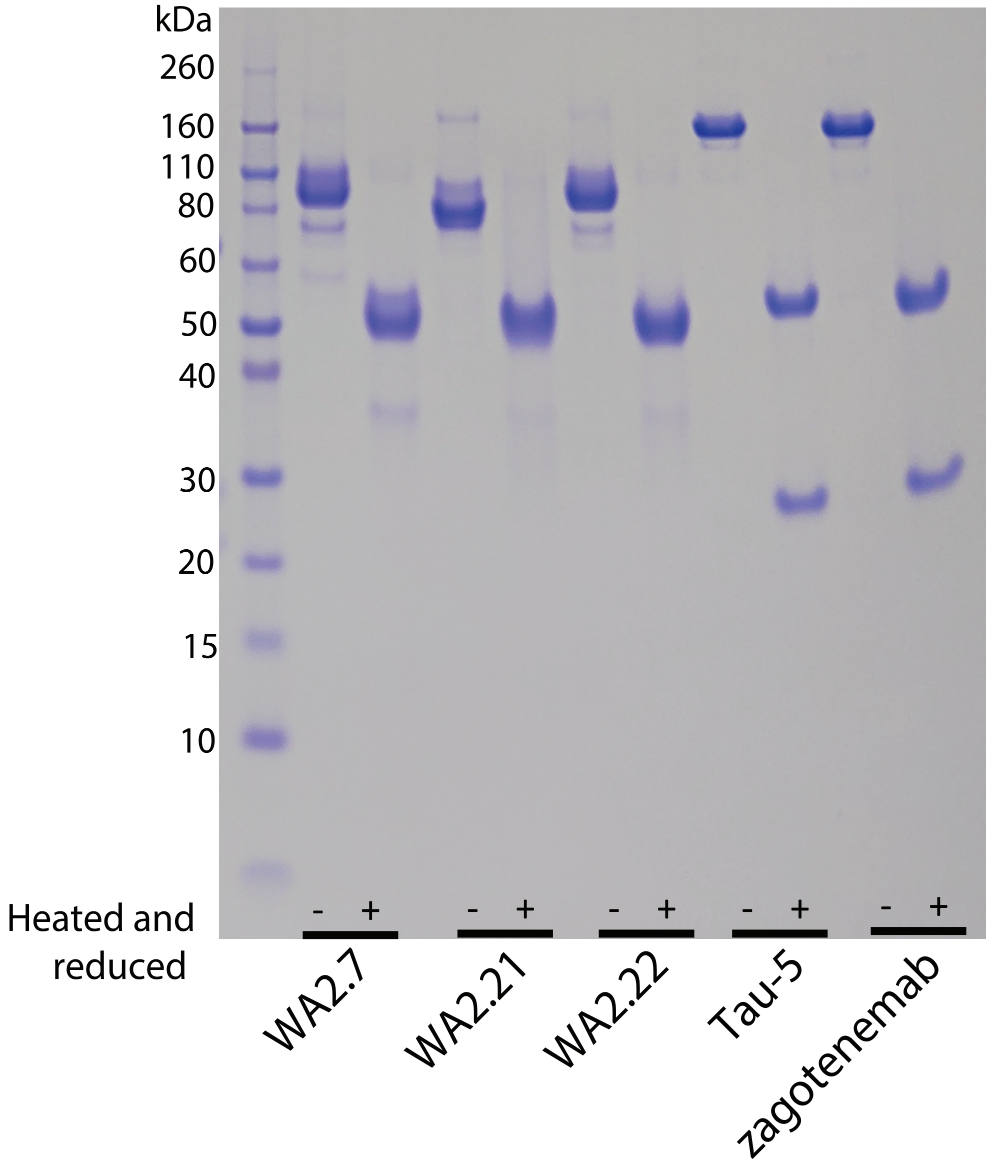


**Supplemental Figure 3. Characterization of tau nanobodies and antibodies by SDS-PAGE.** Nanobody Fc-fusion proteins and antibodies were analyzed by SDS-PAGE either without (-) or with (+) reduction with β-mercaptoethanol (~50 mM) and heating at 100 °C for ~10 min.

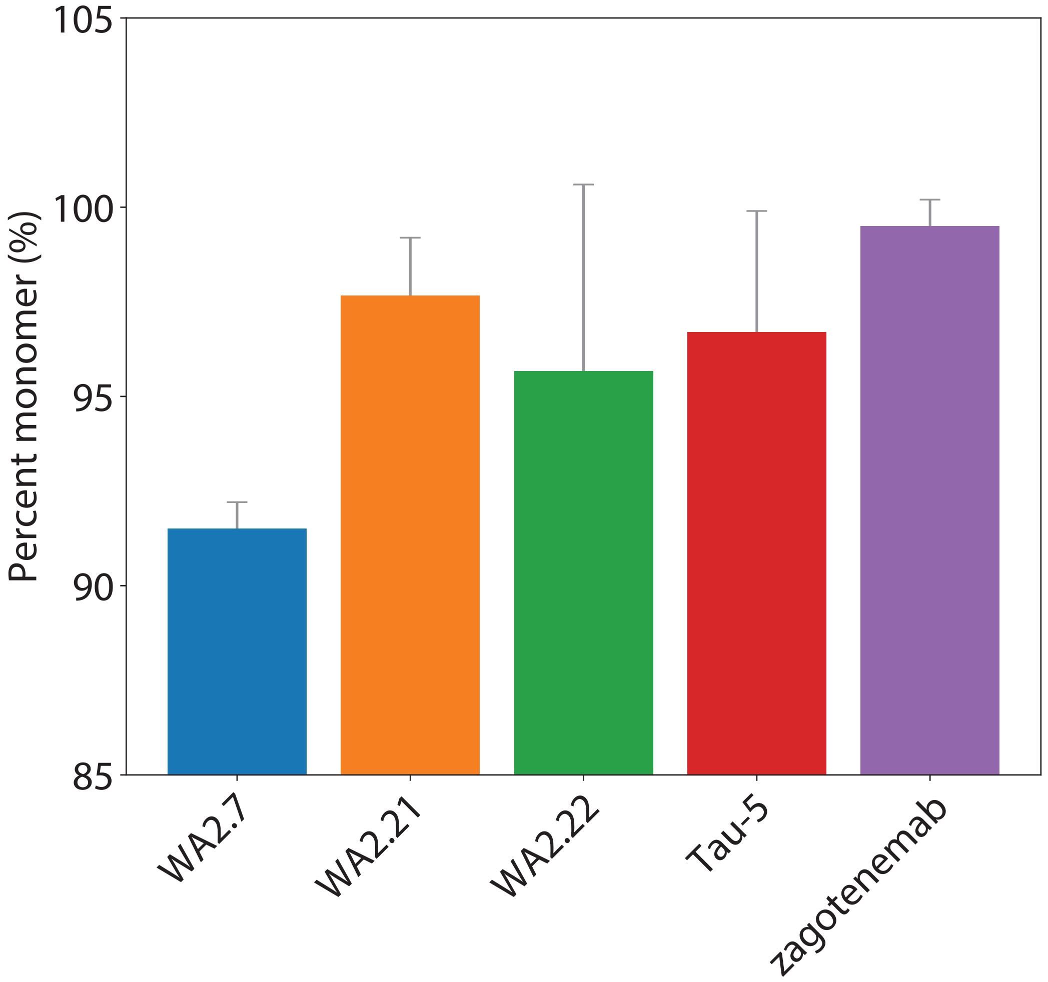


**Supplemental Figure 4. Analytical size-exclusion chromatography analysis of tau nanobodies and antibodies.** The percentage monomer of the tau nanobody Fc-fusion proteins and antibodies was evaluated using analytical size-exclusion chromatography. The data are averages, and the error bars are standard deviations for 2-3 independent experiments.

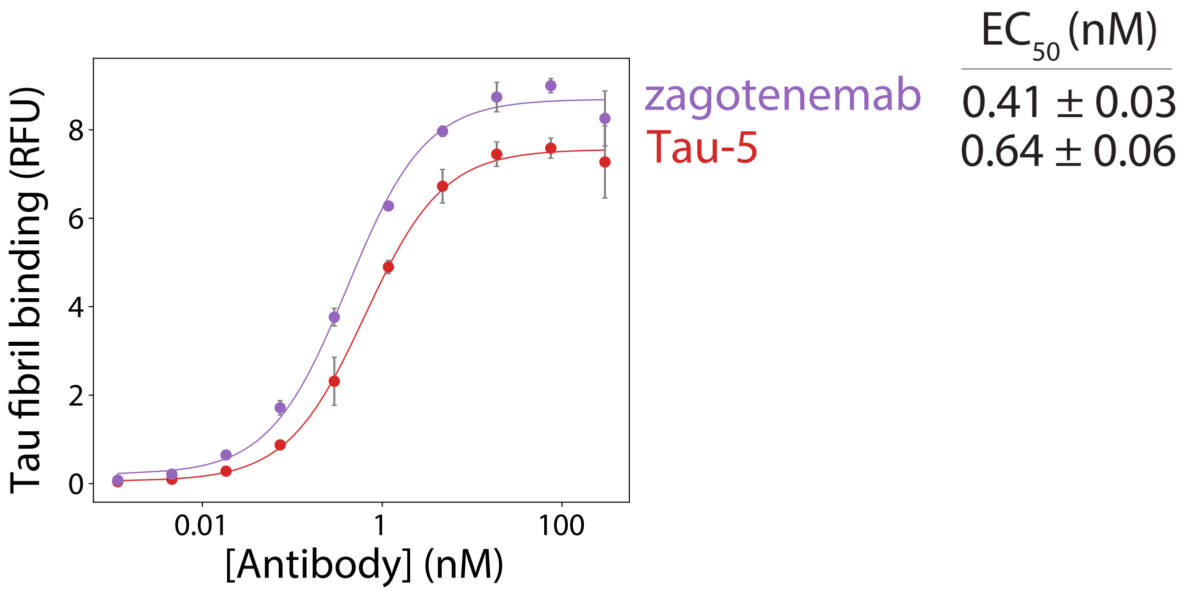


**Supplemental Figure 5. Affinity of control tau antibodies.** The affinities of zagotenemab and Tau-5 were determined using flow cytometry. Various concentrations of antibodies were incubated with tau fibril-coated magnetic beads. Antibody binding was detected using an anti-human Fc 647 secondary antibody. Mean binding signal at each antibody concentration was then determined using flow cytometry. The data are averages, and the errors are standard deviations for three independent experiments.

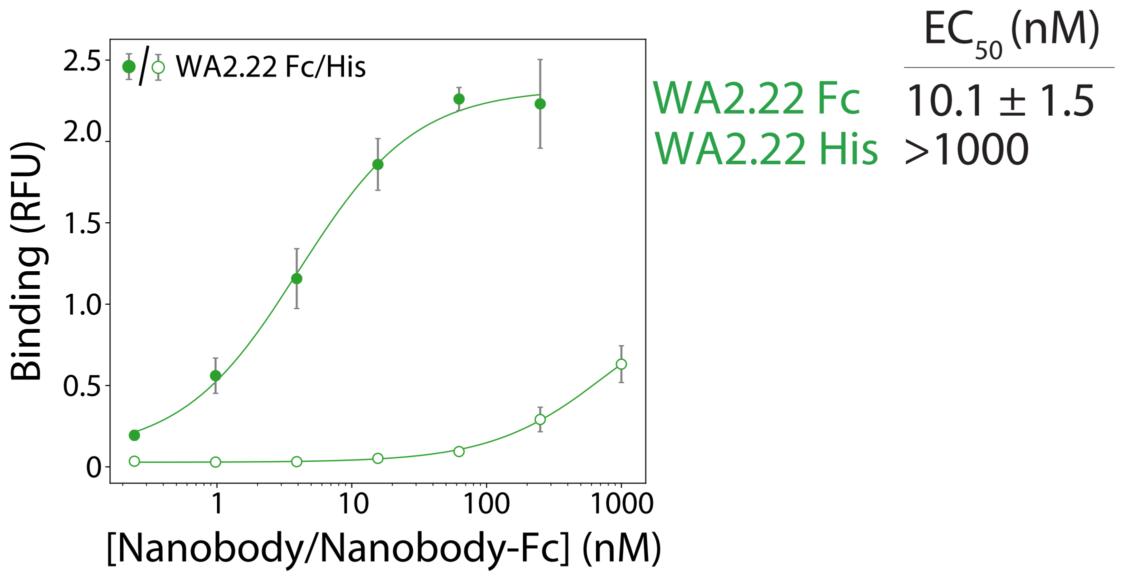


**Supplemental Figure. 6. Comparison of WA2.22 monovalent and bivalent affinities. T**he affinities of WA2.22 as a monovalent nanobody (6x His-tag at C-terminus) and bivalent Fc-fusion protein (6x His- and FLAG-tags at C-terminus) were determined using flow cytometry. WA2.22 or WA2.22 Fc-fusion proteins at various concentrations were incubated with tau fibril-coated magnetic beads. Binding signal was detected using a polyclonal anti-His tag primary antibody and a fluorescently-labeled secondary antibody (Alexa Fluor 647). The mean binding signal at each concentration of WA2.22 or WA2.22 Fc-fusion protein was then determined using flow cytometry. The data are averages, and the error bars are standard deviations for two independent experiments.

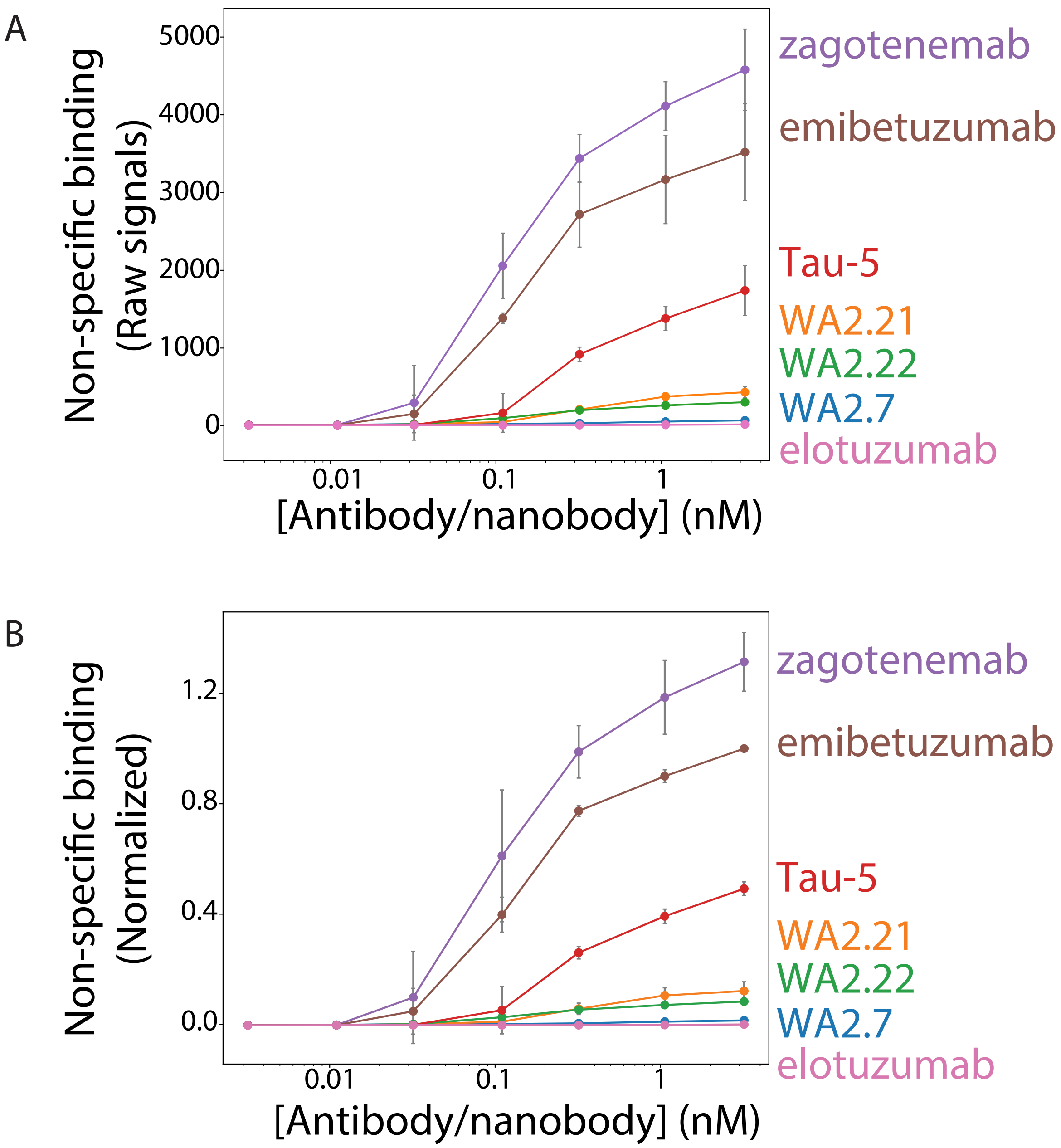


**Supplemental Figure 7.** Comparison of normalized and raw measurements of non-specific binding for tau nanobodies and antibodies. (A) The raw values and (B) normalized values (as shown in Figure 7) obtained for non-specific binding are reported. Normalization was performed by setting the value of non-specific binding at the highest antibody concentrations to 1 for emibetuzumab (high non-specific binding) and 0 for elotuzumab (low non-specific binding), and scaling all other values accordingly. The data are averages, and the error bars are standard deviations for three independent experiments.
